# Combining machine learning and publicly available aerial data (NAIP and NEON) to achieve high-resolution remote sensing of grass-shrub-tree mosaics in the Central Great Plains (U.S.A.)

**DOI:** 10.1101/2025.02.16.638503

**Authors:** Brynn Noble, Zak Ratajczak

## Abstract

Woody plant encroachment (WPE)—a phenomenon similar to species invasion—is shifting many grasslands and savannas into shrub and evergreen-dominated ecosystems. Tracking WPE is difficult because shrubs and small trees are much smaller than the coarse resolution of common remote sensing platforms (> 10 m^2^) and the impassibility of encroaching woody thickets slows ground-based approaches. Many agencies have been investing in fine resolution (< 2 m^2^) remote sensing through programs such as the United States Department of Agriculture (USDA) National Agriculture Imagery Program (NAIP) and the National Ecological Observatory Network (NEON). Both use low-flying planes and provide data to end users in an easy-to-use format at large spatial extents. By removing entry barriers, these publicly available open-source programs could increase the accessibility and extent of remote sensing. We compared two common methods of machine learning classification of land cover (random forests and support vector machines) factorially crossed with these two freely available remote sensing platforms to determine if we could quickly and accurately develop remote sensing of major vegetation types in a tallgrass prairie landscape undergoing encroachment by shrubs and trees. Our work took place at Konza Prairie Biological Station—a landscape scale experiment that results in a wide range of land cover types. All models had very high overall classification accuracy (>90%), with the NEON-based models a few percent more accurate than NAIP. A model using both inputs had the highest accuracy. However, the accuracies of NAIP and NEON models differed for woody vegetation: compared to NEON, NAIP accuracy was, 82-93% compared to 94-98% for shrubs, 72-92% compared to 93-98% for deciduous trees, and 52-78% compared to 83-86% for evergreen trees (specifically *Juniperus virginiana*). NEON-based models relied on canopy height (LiDAR) to make classifications, whereas the several bands of light make similar contributions to accuracy in the NAIP models. Finally, we found that both machine learning approaches had similar accuracy, but random forests ran substantially faster. We conclude that with large training datasets, publicly available aerial imagery and similar products (e.g., UAVs, micro-satellites) can produce fine-scale, high-accuracy remote sensing of WPE in this region with low up-front costs.

## Introduction

Woody encroachment, or woody plant expansion (WPE) in grasslands, is negatively affecting grasslands around the world (Briggs et al. 2002, Brandt et al. 2013, Moser et al. 2013, Twidwell et al. 2013, Galgamuwa et al. 2020). In the Great Plains, tallgrass Prairie is undergoing transitions to shrublands of sumacs (*Rhus spp.*), prunus species (*Prunus spp.*), and Rough Leaf Dogwood (Cornus drummondii) and woodlands of deciduous trees and Eastern Red Cedar (*Juniperus virginiana*; henceforth, ERC) (Briggs et al. 2005, Engle et al. 2008, Ratajczak et al. 2014, Meneguzzo and Liknes 2015). In tallgrass prairie, moderate fire exclusion (fire returns around 3 to 10 years) can result in shrubland expansion in just 10 to 30 years (Ratajczak et al. 2014). With near-complete fire exclusion, ERC can expand, becoming a closed canopy in 20 to 40 years (Briggs et al. 2002). In the Flint Hills, the largest remaining landscape of tallgrass prairie (Scholtz and Twidwell 2022), at least 45% of current grasslands are burned so infrequently that they are likely to transition to shrublands or woodlands in the next ten to thirty years unless management practices change (Ratajczak et al. 2016). As WPE progresses, many grassland obligate species could decline further, including Monarch butterflies (Swengel 1996), native bees (Lettow et al. 2018), songbirds (Engle Silber et al. 2024), upland birds including the threatened Lesser Prairie Chicken (Lautenbach et al. 2017), and others (Albrecht et al. 2016). Woody encroachment can also have economic impacts by reducing freshwater recharge (Keen et al. 2022) and forage for commercial grazers (Morford et al. 2022).

One challenge of studying woody plant encroachment is logistical difficulties. Woody encroachment creates thickets of dense and often thorny vegetation, which can be challenging to measure, pass through, and in extreme cases, measuring the vegetation alters the vegetation itself. Remote sensing could allow grassland ecologists to study woody encroachment more accurately, quickly, and extensively than is possible with on-the-ground approaches. In instances where historical ground-based measurements are unavailable, remote sensing can also provide historical context when aerial imagery is available. This study aimed to determine what combinations of machine learning approaches and data sources produce the most accurate and fastest combination for remote sensing of woody plant encroachment.

Remote sensing and machine learning have long been used to classify land use and land cover (e.g., Laliberte et al. 2004, Asner et al. 2012, Brandt et al. 2013, Allred et al. 2021). Until recently, such studies often used extensive but coarse resolution satellite data—such as the 10 m to 50 m resolution of longer-running satellites (Allred et al. 2021, Galgamuwa et al. 2020, Kranjčić et al. 2019, Nguyen et al. 2020, Thanh Noi and Kappas 2018, Frietag et al. 2021).

Woody encroachment can be difficult to track with coarse-resolution imagery because shrubs and smaller trees are much smaller than the minimum grain size of common satellite data (e.g., <10 m^2^; Whiteman and Brown 1998, Brandt et al. 2020). The growing availability of higher resolution remote sensing from uncrewed aerial vehicles (UAVs), low-flying planes, and high-resolution micro-satellites could rapidly improve our ability to remote sense shrubs and other forms of woody encroachment (Toth and Jóźków 2016, Maxwell et al. 2017, Nagy et al. 2021, Soubry and Guo 2022). For instance, high-resolution remote sensing allowed Brandt and colleagues (2020) to identify millions of small trees across Northern Africa—a region thought to house few trees, because more coarse-resolution resolution data products could not identify the smaller trees of this region.

There are multiple options for remote sensing vegetation. Here we focus primarily on publicly available data from low-flying planes, which combine high resolution with a low upfront costs to the end user and large spatial extent. In the United States, the United States Department of Agriculture (USDA) National Agriculture Imagery Program (NAIP) has provided the most consistent, widespread, and freely available high-resolution remote sensing data in the United States. This product was a large investment, with mixed impact on peer-reviewed literature (reviewed by Maxwell et al. 2017). Recent investments from the U.S.A. National Science Foundation now provide high-resolution remote sensing data through the National Ecological Observatory Network’s (NEON) aerial observation platform (reviewed by Nagy et al. 2021). NEON covers a smaller spatial extent than NAIP (81 total sites versus the entire continental U.S.A.), but provides a much wider range of data per site, including LiDAR, hyperspectral data, and derived products. These additions are promising, but because their adoption by ecologists is still less common than many expected (e.g., Maxwell et al. 2017). Therefore, several questions remain: is the learning curve of using these products preventing widespread use? How accurate are NEON’s derived products? For instance, at one site, a recent study found that some derived NEON products had a weak correlation with ground-based measurements (Pau et al. 2022). On the other hand, there have been successful uses of NEON to identify tree crowns (Weinstein et al. 2024). Thus, we are left with a mixed assessment—high-resolution publicly available imagery shows promise for detecting WPE, but use of these platforms is still relatively rare compared to UAVs and satellites.

Here, we assess the accuracy of using high-resolution remote sensing to measure woody plant encroachment by three functional types of woody plants—shrubs, deciduous trees, and ERC. We assessed the value-added of NEON for remote sensing woody plant encroachment, over the more widely available and longer-running NAIP program. For instance, more data typically increases accuracy, but inputs such as hyperspectral data are data-heavy, require substantial computing power, and are less accessible to many users. We aimed to determine the input and methods necessary to accurately measure WPE while limiting computational effort and using data products available to a reasonably skilled graduate student, post-doc, professor, or user at a private, state, or federal agency. Therefore, we restricted our use of NEON data to “off the shelf” data products, such as NEON’s data product “estimated canopy height” based on LiDAR and vegetation indices, including the widely used Normalized Difference Vegetation Index (NDVI) and more specific indices such as foliar nitrogen and total lignin. Our decision was motivated by observations that a lack of certain computational skills might impede the wider usage of NEON remote sensing products (see Maxwell et al. 2017 and Nagy et al. 2021 for reviews).

We performed a factorial modeling exercise exploring the role of more remote sensing inputs (NAIP RGB and Near-Infrared, NEON vegetation indices and LiDAR), crossed with model sophistication, comparing support vector machines (SVMs) and random forests models (RFs). We hypothesized that: 1) NEON would increase accuracy, primarily due to the addition of canopy height estimated using LiDAR; 2) shrubs would have lower accuracy than grasses and trees, because shrub height and traits fall in between herbaceous species and trees, and because there is a high diversity of traits within the shrub community (Wedel et al. 2025); and 3) ERC would have high accuracy, due to its unique evergreen leaf type compared to the other functional groups we considered.

## Methods

### Site Description

Konza Prairie Biological Station (KPBS), is a National Science Foundation long-term ecological research (LTER) site with 3,487 ha of native unplowed tallgrass prairie located in the Flint Hills in northeastern Kansas (Fig. 1). KPBS has high seasonal variability with an average high of 26.6⁰ C in July and –2.7⁰ C in January. The annual rainfall is 835 mm, with 75% falling during the growing season. The soils are non-glaciated with thin rocky upland soils (mainly from the Florence series), deeper lowlands (often from the Tully series), and complex benches, outcrops, and slopes that connect these two soil types.

**Figure 1:**
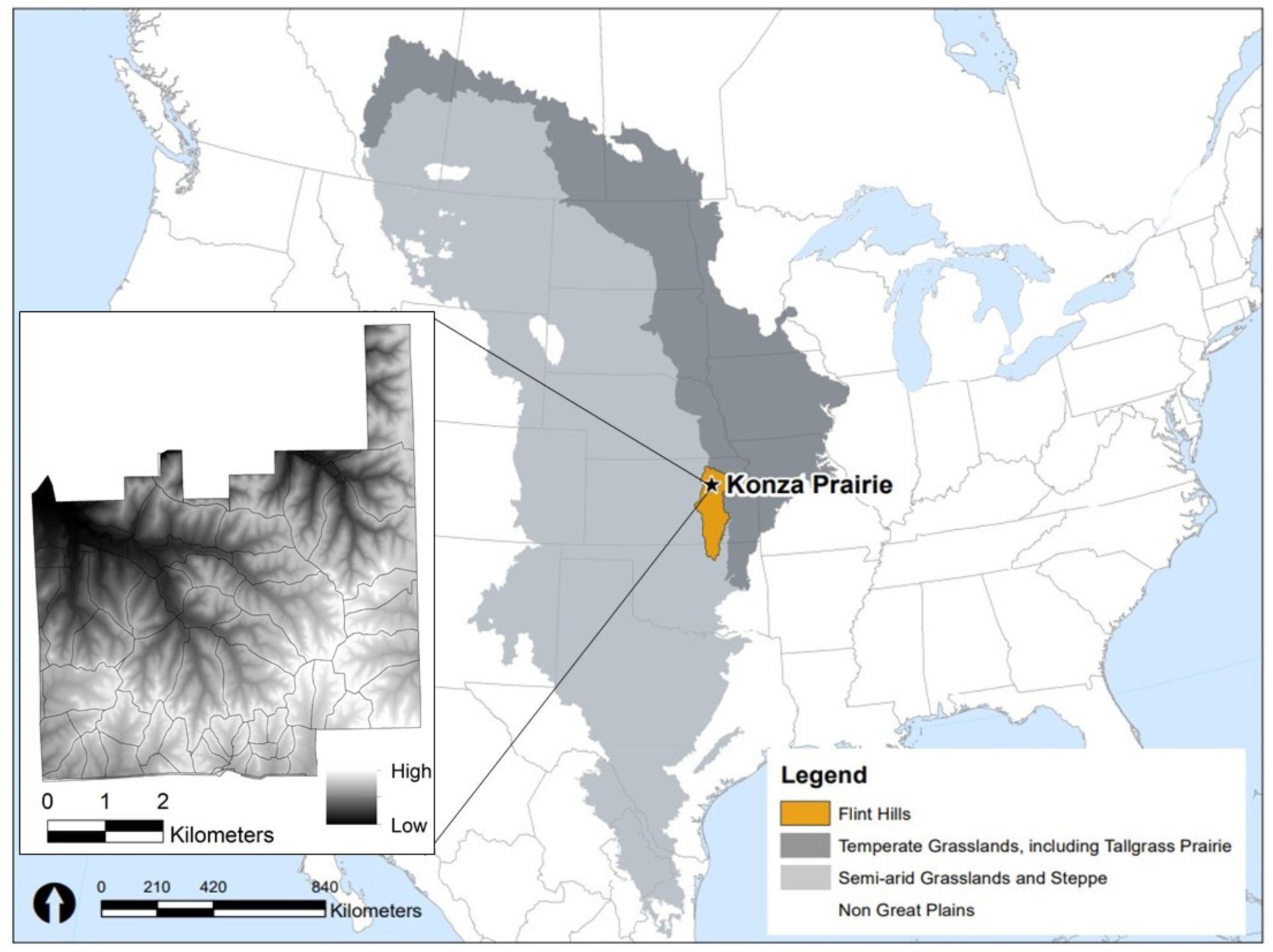
An estimate of the historical extent of arid and semi-arid Great Plains grasslands (light grey; based on EPA ecoregions), temperate Great Plains grasslands (dark grey; based on EPA ecoregions), the Flint Hills ecoregion (orange), and our study site (black star). The map inset shows an elevation map of our study site, Konza Prairie Biological Station.

KPBS consists of 60 different management units, with replicates spanning 1-, 2-, 3-, 4-, and 20-year fire frequencies, as well as ungrazed, grazed by bison, or grazed by cattle. These different treatments have created a mosaic of contrasting land covers, including areas dominated by herbaceous species, shrubs, deciduous trees, and evergreen trees. The herbaceous-dominated areas can be floristically diverse, with high grass dominance in areas without bison or cattle, and mosaics of tallgrasses, short-grass grazing lawns, and patches of tall forbs in areas with bison and cattle (Ratajczak et al. 2022). Areas dominated by shrubs typically have little to no herbaceous species (Briggs et al. 2002, Ratajczak et al. 2011, Ratajczak et al. 2014) and dominant shrubs (primarily the species *Cornus drummondii*) are all clonal, creating “islands” of ramets that can reach over 10 m in diameter. Shrub heights range from <0.5 m for young clonal stems to over 3 m for older stems (Tooley et al. 2024). At the lowest elevations and some intermittent streams, a full riparian forest has become established, dominated primarily by oaks (mostly Chinqapin oak, *Quercus muehlenbergii* and Burr oak, *Quercus macrocarpa*). Outside of these lowlands, the height and continuity of deciduous trees is lower, with species including honeylocust (*Gleditsia triacanthos*), redbud (*Cercis canadensis*), and several elm species. The only evergreen trees known to occur on site is ERC (*Juniperus virginiana*), primarily in areas without bison and frequent fire (Noble, Noble, and Ratajczak, *In review*).

### Imagery

We used two remote sensing platforms: USDA NAIP and NSF NEON. Each data source was tested alone and together (NAIP+NEON) for a total of three models for each machine learning method, resulting in six models. Table 1 outlines the inputs used for each image. The images were captured in separate years, but between these two years there were no major weather fluctuations (Ratajczak et al. 2022) nor major fires. Therefore, significant changes in vegetation between these two time periods is unlikely.

**Table 1:** Summary of input variables and their descriptions used from each source.

| Source | # bands | Inputs | Input Description |
| --- | --- | --- | --- |
| USDA<br>NAIP | 9 | Red | Redness of each pixel |
|  |  | Green | Greenness of each pixel |
|  |  | Blue | Blueness of each pixel |
|  |  | Red Neighborhood* | Avg. redness of surrounding pixels |
|  |  | Green Neighborhood* | Avg. greenness of surrounding pixels |
|  |  | Blue Neighborhood* | Avg. blueness of surrounding pixels |
|  |  | Near-infrared | Value of near-infrared wavelength |
|  |  | Near-infrared<br>Neighborhood* | Avg. values of near-infrared of surrounding pixels |
|  |  | NDVI | Calculated from NIR and red bands, indicates live green<br>vegetation density |
| NSF<br>NEON | 8 | Enhanced vegetation<br>index (EVI) | Similar to NDVI, estimates vegetation greenness and<br>biomass |
|  |  | Normalized difference<br>nitrogen index (NDNI) | Relative nitrogen concentration in canopy |
|  |  | Normalized difference<br>lignin index (NDLI) | Uses shortwave IR to estimate lignin content in canopy |
|  |  | Soil-adjusted vegetation<br>index (SAVI) | Reduces soil brightness in areas where vegetation cover is<br>low |
|  |  | Atmospherically resistant<br>vegetation index (ARVI) | Reduces atmospheric noise from dust, smoke, rain, etc. |
|  |  | NDVI | Calculated from NIR and red bands, indicates live green<br>vegetation density |
|  |  | NDVI neighborhood* | Avg. NDVI values of surrounding pixels |
|  |  | Canopy height (LiDAR) | Height of canopy above bare earth |
| NEON +<br>NAIP | 17 | All of the above |  |

\* “Neighborhood” refers to an average of all neighboring pixels using the Queen’s rule to determine which pixels are included in as neighbors

*NAIP imagery*: This image was sourced from the USDA NAIP (USDA 2019a,b, Noble and Ratajczak 2022 for the final imagery used). The image was captured on July 10, 2019 by low-flying planes and has a native resolution of 0.6 m^2^. We used bilinear interpolation to transform each image to 2 m^2^ pixels snapped to a common grid with all other images/inputs.

The image sourced from NAIP contains nine bands. The native imagery provide four bands (red, green, blue, infrared), and we derived five more (red neighborhood, green neighborhood, blue neighborhood, infrared neighborhood, and NDVI; Table 1). Neighborhood calculations take the average value of all surrounding pixels. The red neighborhood calculation, for example, averages the redness of the pixels immediately surrounding each pixel, including the diagonal neighbors (known as the ‘Queen’s rule’). We added neighborhood averages after early rounds of machine learning erroneously misclassified some deep as ERC.

*NEON imagery:* NSF NEON imagery was captured in June 2020 by low-flying airplanes and has a resolution of 1 m^2^, which was upscaled to the same resolution and grid as NAIP using bilinear interpolation (NEON 2020a,b, accessible at Noble and Ratajczak 2022). NEON provides many data products meant to capture biophysical attributes of vegetation. For example, NDVI is derived from dividing the difference between near-infrared (NIR) and red bands by adding NIR and red bands. We used eight derived vegetation indices from NEON: enhanced vegetation index (EVI), normalized difference nitrogen index (NDNI), normalized difference lignin index (NDLI), soil-adjusted vegetation index (SAVI), atmospherically resistant vegetation index (ARVI), NDVI, NDVI neighborhood, and canopy height (LiDAR; Table 1). Leaf area index was not included because it is calculated based on another index, SAVI. NEON’s 10-cm RGB was unusable for machine learning due to distortions along seamlines, however these distortions were not in the derived products.

### Machine Learning Methods

Machine learning uses a set of user inputs (training data) to learn and classify unknown data. We compared two different supervised machine learning methods for this project, SVM and RF, as these are the most common methods used in remote sensing today (Thanh Noi and Kappas 2018, Sheykhmousa et al. 2020).

*Support Vector Machines:* Support Vector Machines (SVM) are a supervised nonparametric classification technique that use a fixed optimal hyperplane to split the data into the desired number of discrete categories (Burges 1998). In the simplest form, a linear line separates two-dimensional data into two categories (Mountrakis et al. 2011; Fig. 2), but SVMs are popular for their ability to work with high-dimensional data (Sheykhmousa et al. 2020).

**Figure 2:**
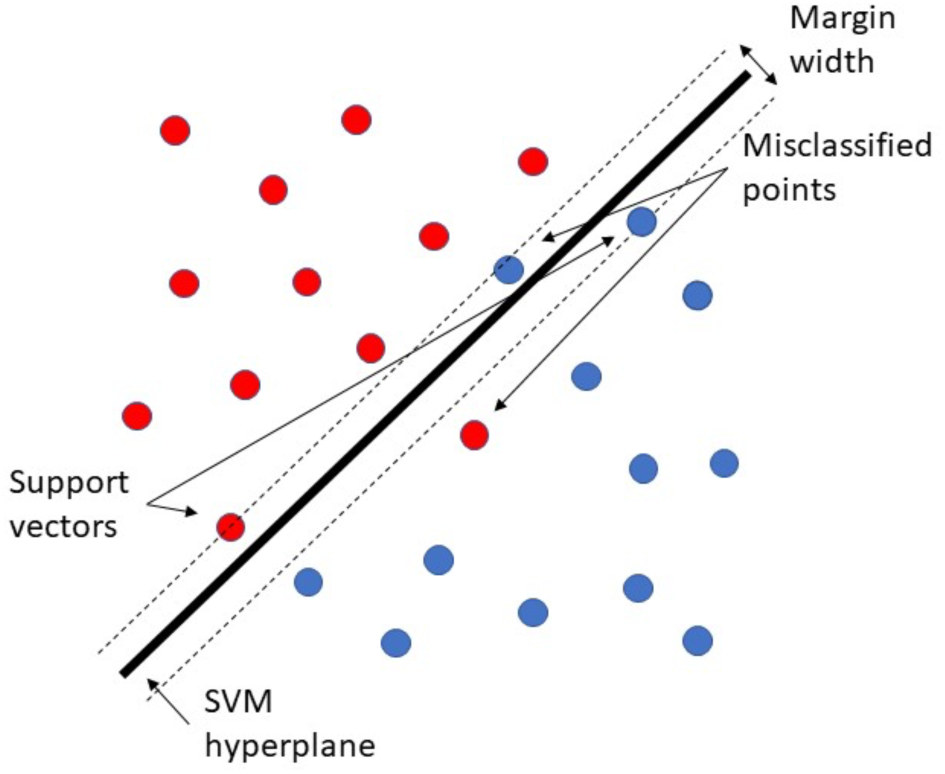
Simple linear form of SVM. Adopted from Burges 1998, which is the type of SVM used in this study.

Each pixel of a remote sensing image is a series of numbers, with one value from each input, which are mapped with each input as a new dimension during SVM model creation. Points that lie closest to the hyperplane are support vectors and are the most important in determining the decision boundary. While many linear hyperplanes may exist in the data, SVM chooses the largest margin between points, allowing for some misclassification. In real-life applications, many datasets do not have a linear break between data. SVMs can easily circumvent this problem by shifting the data into a higher dimension using kernels, which separates the data even further to allow for a linear hyperplane to split the data. Furthermore, SVMs are often used in the remote sensing field for their accuracy even with small training sets (Thanh Noi and Kappas 2018, Mantero et al. 2005), making them a strong choice for large study areas.

SVMs were run in program R (v4.0.5; R Core Team 2021) using the ‘e1071’ package, and model parameters were optimized using the ‘best.svm’ function (v1.7-6; Mayer et al. 2021; see Table S1 for final model parameters).

*Random Forests:* RFs are a nonparametric supervised machine learning technique, with the advantage of being less prone to model over-fitting (e.g., low out-of-bag error) and being able to give estimates of which remote sensing inputs are the most important for model accuracy. The building block of RF are decision trees, which use nodes to split data into smaller and smaller subsets to predict the pixel class. RFs use a bagging approach when building trees, where each decision tree is built with a random selection of input variables, creating a forest of different tree structures to limit overfitting (Evans et al. 2011). The user can set the number of decision trees for each model (ntree) and the number of input variables used to split each node (mtry), but many users rely on the default values of ntree (500) and mtry (square-root of number of inputs; Thanh Noi and Kappas 2018). RFs make predictions based on majority voting; each decision tree outputs a predicted class and the class which is predicted the most times in the forest is the overall predicted classification (Sheykhmousa et al. 2020). For example, if a RF has 100 trees, and 56 tress predict a pixel as grassland, 33 trees predict that the pixel is shrubland, and 11 trees predict deciduous tree, then the ensemble RF model would predict that pixel to be grassland.

Adding decision trees can improve accuracy, but it will increase the model run time and required computing capacity. Models with an excessive number of trees yield diminishing returns in accuracy. Adding trees also runs the risk of model over-fitting. Lastly, RF models can determine the GINI decrease for each input, which measures the amount of accuracy lost by an input variable, indicating the relative importance of each.

RF models were run in program R (v4.0.5; R Core Team 2021) using the ‘randomForest’ package and model inputs were optimized using the ‘best.randomForest’ function (v4.6.14; Liaw and Wiener 2002). The NAIP+NEON model had an ntree of 500 (the number of trees in the rando forest model) and mtry of 4 (the number of variables assessed at each node of each decision tree). Both single-source models (NAIP and NEON) had an ntree of 500 and mtry of 3.

### Training Data

We considered five land cover categories: (1) grassland (herbaceous-dominated areas); (2) shrubs; (3) deciduous trees; (4) ERC trees; (5) and other (roads, water, and buildings). We created training through a combination of ground-truth polygons collected using high-precision GPS units (with below 2 m error) and computer-drawn polygons. Ground-truth polygons data collection occurred from June to August 2021 and were collected using semi-stratified random sampling. We created computer-drawn by tracing vegetation types, based on a 2020 NEON RGB-10 cm imagery and publicly available 1 m^2^ RGB (a 2019 image from Maxar technologies available at Google Earth). Neither of these images were used in the SVMs or RFs. We drew polygons in locations where species were known from time in the field or where were well-known (in particular, buildings, water, and roads). We collected 3,635 training polygons, with training proportions of each cover class reflecting landscape cover, resulting in 300,328 2 x 2 m pixels of known vegetation type, totaling 3.42% of the total study area (Table 2). We used 70% of the polygons to train the models and we held out the remaining 30% for evaluation of model accuracy. Polygons were split into training and evaluation groups before being pixelized to reduce inflated accuracies due to spatial autocorrelation. After we converted polygons into pixels, the proportion of training and evaluation pixels remained similar (71% and 29%, respectively).

**Table 2:** Summary of training data. Training proportions are similar to landscape cover.

| Class | # Ground-truthed polygons | # Computer-drawn polygons | Total polygons | Total Pixels | Total area (m<sup>2</sup>) | % of total training |
| --- | --- | --- | --- | --- | --- | --- |
| Deciduous Trees | 68 | 620 | 688 | 37,215 | 150,533.5 | 12.7% |
| Grass | 246 | 160 | 406 | 179,799 | 719,013.9 | 60.3% |
| Easter Red Cedar | 51 | 506 | 557 | 5,537 | 22,751.6 | 1.9% |
| Shrubs | 341 | 1578 | 1919 | 71,101 | 285,690.1 | 24% |
| Other* | 0 | 65 | 65 | 6,676 | 13,484.3 | 1.1% |
| Total | 706 | 2929 | 3635 | 300,328 | 1,191,473 | 100% |

**\*Other = water, roads, and buildings**

### Assessing accuracy

When assessing the accuracy of each model, four aspects are considered: producer accuracy (PA), user accuracy (UA), overall accuracy (OA), and Kappa. PA refers to the number of pixels correctly classified from the evaluation data. In other words, the proportion of ‘grassland’ pixels in the evaluation data also appear as ‘grassland’ in the final classified image. UA refers to the number of pixels which classified as, for instance, grassland that were actually grassland on the ground (i.e., in the evaluation data). OA is an estimate of accuracy among all predicted cover types and is a ratio of the total number of correctly classified pixels to the total number of pixels. Lastly, the Kappa coefficient measures the accuracies by comparing the classification outcome versus randomly assigning values. Kappa ranges from –1 to 1, with 0 indicating that the model performed on par with randomly assigning values, <0 indicating that the model performed worse than random, and >0 indicating that the model performed better than random.

### Run time and other logistics

Run time can become a consideration for some machine learning approaches, requiring PIs and/or students to learn new techniques to complete more computationally intensive tasks. We report run time for both model training and extrapolation to the remainder of our site (a total of 8,781,520 pixels), to give an estimate of trade-offs between model accuracy versus computational efficiency. For reference, these models were run on a Dell XPS 8930, with relevant specifications of 64 GB RAM and an Intel^®^ Core™ i9-9900K processor, with 3.6 GHz speed, eight cores, and the ability to perform 16 threads. Note that program R runs most processes through the RAM.

## Results

### Model accuracy overview

Our two modelling approaches (SVMs vs RFs) yielded nearly identical OA (<2.6% difference; Table 3). NEON models generally performed better than NAIP, and the NAIP+NEON models were more accurate than any single-source model. NAIP+NEON had almost no difference between RFs (OA: 97.7%, Kappa: 0.962) and SVMs (OA: 97.7%, Kappa: 0.964; Table 3). NEON also had very little difference in accuracies between RF (OA: 97.2%, Kappa: 0.951) and SVM (OA: 96.8%, Kappa: 0.945; Table 3). NAIP had the largest difference between classification method, with RFs slightly more accurate (OA: 92.9%, Kappa: 0.877) than SVMs (OA: 90.3%, Kappa: 0.831; Table 3). Classified maps comparing all three data sources are shown in Fig. 3d-e, where the machine learning method was RFs for all panels.

**Figure 3:**
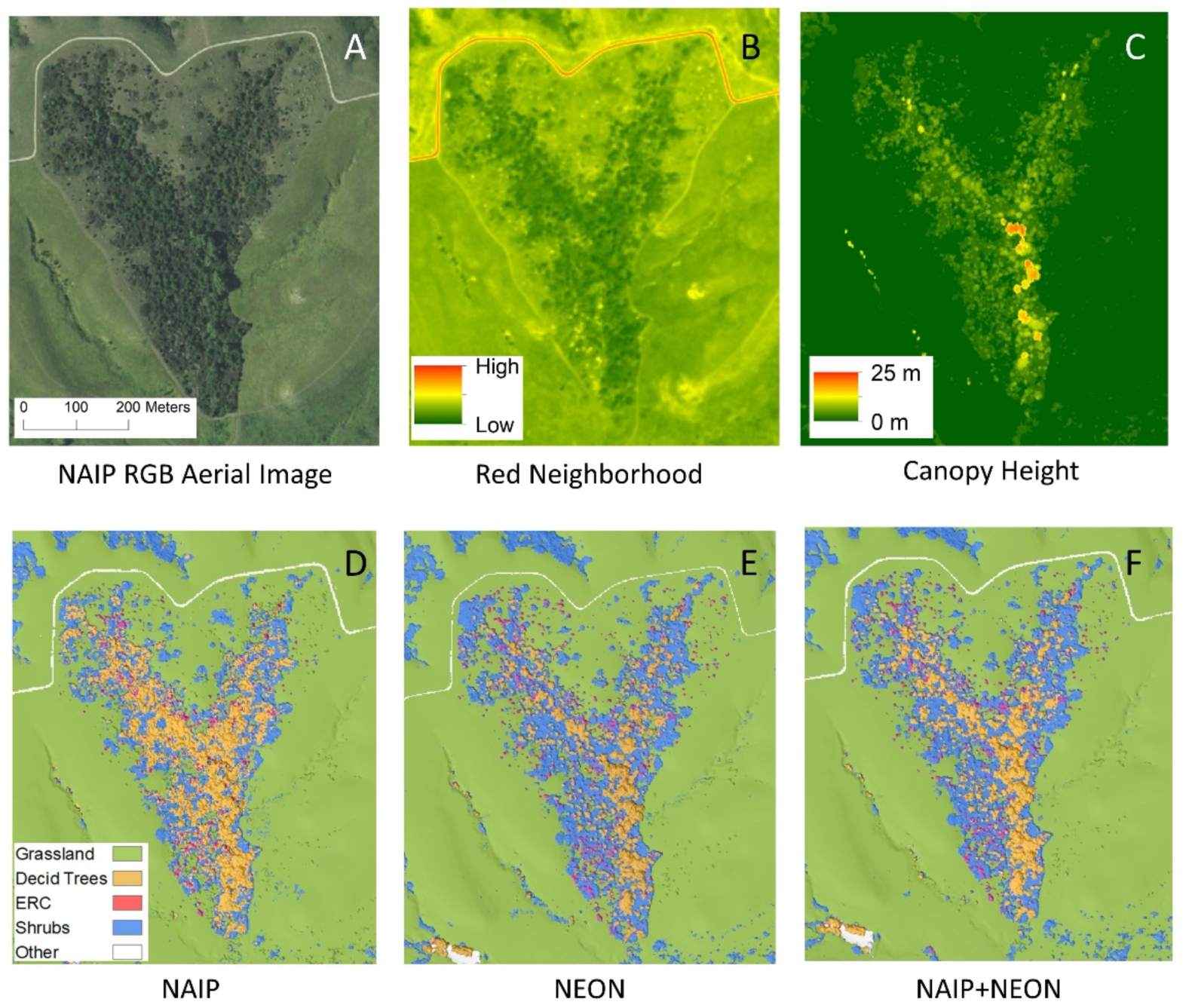
A small portion of the study area, zoomed in to show detail: A) Aerial imagery from NAIP RGB (red-green-blue; naked eye view); B) visual of values in most important NAIP input for model training; C) visual of values most important NEON input for model training; D-F) visuals of RF classified models for all three images.

**Table 3:** Accuracy results of all image and machine learning methods.

| Source | Machine learning method | OA* | Kappa | Accuracy | Deciduous Trees | Grassland | ERC | Shrubs | Other** |
| --- | --- | --- | --- | --- | --- | --- | --- | --- | --- |
| NAIP | SVM | 0.903 | 0.831 | PA | 0.72 | 0.98 | 0.52 | 0.85 | 0.94 |
|  |  |  |  | UA | 0.89 | 0.94 | 0.78 | 0.82 | 0.97 |
|  | RF | 0.929 | 0.877 | PA | 0.77 | 0.98 | 0.56 | 0.93 | 0.96 |
|  |  |  |  | UA | 0.92 | 0.97 | 0.75 | 0.85 | 0.97 |
| NEON | SVM | 0.968 | 0.945 | PA | 0.96 | 0.99 | 0.84 | 0.98 | 0.97 |
|  |  |  |  | UA | 0.98 | 0.98 | 0.86 | 0.94 | 0.97 |
|  | RF | 0.972 | 0.951 | PA | 0.93 | 0.99 | 0.83 | 0.98 | 0.96 |
|  |  |  |  | UA | 0.98 | 0.99 | 0.85 | 0.94 | 0.97 |
| NAIP + NEON | SVM | 0.977 | 0.964 | PA | 0.95 | 0.99 | 0.87 | 0.97 | 0.97 |
|  |  |  |  | UA | 0.97 | 0.98 | 0.90 | 0.96 | 0.99 |
|  | RF | 0.977 | 0.962 | PA | 0.94 | 0.99 | 0.85 | 0.98 | 0.96 |
|  |  |  |  | UA | 0.98 | 0.99 | 0.87 | 0.95 | 0.98 |

\*OA: overall accuracy; PA: producer accuracy; UA: user accuracy.
\*\*Other = water, roads, and buildings

For all models, the grassland and “other” categories had values of 94% or above for both UA and PA for all combinations of machine learning methods and data sources (Table 3). The three categories of woody plants (shrubs, deciduous trees, and ERC) were more difficult to classify, but still had high accuracy ratings. Shrubs and deciduous trees had very high PA and UA accuracies (>92%) in NEON and NAIP+NEON, but NAIP alone performed slightly worse, with PA of 85-93% for shrubs and 72-77% for deciduous trees, and UA of 82-85% for shrubs and 89-92% for deciduous trees (Table 3). Deciduous trees were most often misclassified as shrubs, and shrubs were most often misclassified as deciduous trees (Tables 4-9). All models had the lowest accuracies for ERC, but NAIP performed particularly poorly. PA and UA for NAIP-based ERC were both low, but PA was especially low at 52-56%, compared to 75-78% UA (Tables 3-5). This low PA means that the models are undercounting ERC by classifying it as another cover class (mostly deciduous trees), rather than misclassifying other categories as ERC (also referred to as an error of commission)(Tables 4 & 5).

**Table 4:** Confusion matrix for SVM NAIP; columns represent class of evaluation pixels, and rows represent class of model classified pixels.

| **SVM NAIP** | **Deciduous Trees** | **Grassland** | **Eastern Red Cedar** | **Shrub** | **Other** | **User accuracy** |
| --- | --- | --- | --- | --- | --- | --- |
| **Deciduous Trees** | 9303 (72.4%) | 19 (0%) | 457 (28.1%) | 665 (3.2%) | 6 (0.4%) | 0.89 |
| **Grassland** | 325 (2.5%) | 48825 (98.3%) | 163 (10%) | 2437 (11.8%) | 72 (4.8%) | 0.94 |
| **Eastern Red Cedar** | 198 (1.5%) | 22 (0%) | 839 (51.6%) | 18 (0.1%) | 3 (0.2%) | 0.78 |
| **Shrub** | 2999 (23.4%) | 789 (1.6%) | 168 (10.3%) | 17521 (84.9%) | 1 (0.1%) | 0.82 |
| **Other** | 18 (0.1%) | 19 (0%) | 0 (0%) | 0 (0%) | 1405 (94.5%) | 0.97 |
| **Producer accuracy** | 0.72 | 0.98 | 0.52 | 0.85 | 0.94 |  |

\*The percent of pixels for each row by column is given in parentheses to gauge errors of omission and commission

**Table 5:** Confusion matrix for RF NAIP; columns represent class of evaluation pixels, and rows represent class of model classified pixels*.

|  | **Deciduous**<br>**Trees** | **Grassland** | **Eastern Red**<br>**Cedar** | **Shrub** | **Other** | **User**<br>**accuracy** |
| --- | --- | --- | --- | --- | --- | --- |
| **Deciduous**<br>**Trees** | 9852 (76.7%) | 26 (0.1%) | 439 (27%) | 370 (1.8%) | 10 (0.7%) | 0.92 |
| **Grassland** | 305 (2.4%) | 48667 (98%) | 127 (7.8%) | 1010 (4.9%) | 62 (4.2%) | 0.97 |
| **Eastern Red**<br>**Cedar** | 276 (2.1%) | 28 (0.1%) | 914 (56.2%) | 3 (0%) | 3 (0.2%) | 0.75 |
| **Shrub** | 2389 (18.6%) | 927 (1.9%) | 147 (9%) | 19249<br>(93.3%) | 0 (0%) | 0.85 |
| **Other** | 21 (0.2%) | 26 (0.1%) | 0 (0%) | 0 (0%) | 1412 (95%) | 0.97 |
| **Producer**<br>**accuracy** | 0.77 | 0.98 | 0.56 | 0.93 | 0.95 |  |

\*The percent of pixels for each row by column is given in parentheses to gauge errors of omission and commission

**Table 6:** Confusion matrix for SVM NEON; columns represent class of evaluation pixels, and rows represent class of model classified pixels.

|  | **Deciduous Trees** | **Grassland** | **Eastern Red Cedar** | **Shrub** | **Other** | **User accuracy** |
| --- | --- | --- | --- | --- | --- | --- |
| **Deciduous Trees** | 11877 (92.5%) | 15 (0%) | 137 (8.4%) | 114 (0.6%) | 11 (0.7%) | 0.98 |
| **Grassland** | 81 (0.6%) | 49072 (98.8%) | 32 (2%) | 718 (3.5%) | 43 (2.9%) | 0.98 |
| **Eastern Red Cedar** | 193 (1.5%) | 5 (0%) | 1333 (81.9%) | 12 (0.1%) | 1 (0.1%) | 0.86 |
| **Shrub** | 679 (5.3%) | 551 (1.1%) | 125 (7.7%) | 19788 (95.9%) | 1 (0.1%) | 0.94 |
| **Other** | 13 (0.1%) | 31 (0.1%) | 0 (0%) | 0 (0%) | 1431 (96.2%) | 0.97 |
| **Producer accuracy** | 0.92 | 0.99 | 0.82 | 0.96 | 0.96 |  |

\*The percent of pixels for each row by column is given in parentheses to gauge errors of omission and commission

**Table 7:** Confusion matrix for RF NEON; columns represent class of evaluation pixels, and rows represent class of model classified pixels.

|  | Deciduous<br>Trees | Grassland | Eastern Red<br>Cedar | Shrub | Other | User<br>accuracy |
| --- | --- | --- | --- | --- | --- | --- |
| **Deciduous**<br>**Trees** | 11891<br>(92.6%) | 13 (0%) | 161 (9.9%) | 30 (0.1%) | 9 (0.6%) | 0.98 |
| **Grassland** | 83 (0.6%) | 48973<br>(98.6%) | 15 (0.9%) | 420 (2%) | 46 (3.1%) | 0.99 |
| **Eastern Red**<br>**Cedar** | 210 (1.6%) | 8 (0%) | 1346<br>(82.7%) | 17 (0.1%) | 2 (0.1%) | 0.85 |
| **Shrub** | 651 (5.1%) | 642 (1.3%) | 105 (6.5%) | 20165<br>(97.7%) | 2 (0.1%) | 0.94 |
| **Other** | 8 (0.1%) | 38 (0.1%) | 0 (0%) | 0 (0%) | 1428 (96%) | 0.97 |
| **Producer**<br>**accuracy** | 0.92 | 0.99 | 0.83 | 0.98 | 0.96 |  |

\*The percent of pixels for each row by column is given in parentheses to gauge errors of omission and commission

**Table 8:** Confusion matrix for SVM NAIP+NEON; columns represent class of evaluation pixels, and rows represent class of model classified pixels.

| **SVM**<br>**NAIP+NEON** | **Deciduous**<br>**Trees** | **Grassland** | **Eastern**<br>**Red Cedar** | **Shrub** | **Other** | **User**<br>**accuracy** |
| --- | --- | --- | --- | --- | --- | --- |
| **Deciduous**<br>**Trees** | 12157<br>(94.7%) | 34 (0.1%) | 112 (6.9%) | 85 (0.4%) | 12 (0.8%) | 0.98 |
| **Grassland** | 55 (0.4%) | 49388<br>(99.4%) | 23 (1.4%) | 511 (2.5%) | 27 (1.8%) | 0.98 |
| **Eastern Red**<br>**Cedar** | 127 (1%) | 3 (0%) | 1413<br>(86.8%) | 7 (0%) | 7 (0.5%) | 0.90 |
| **Shrub** | 487 (3.8%) | 232 (0.5%) | 79 (4.9%) | 20029<br>(97.1%) | 0 (0%) | 0.96 |
| **Other** | 17 (0.1%) | 17 (0%) | 0 (0%) | 0 (0%) | 1465 (97%) | 0.99 |
| **Producer**<br>**accuracy** | 0.95 | 0.99 | 0.87 | 0.97 | 0.97 |  |

\*The percent of pixels for each row by column is given in parentheses to gauge errors of omission and commission

**Table 9:**
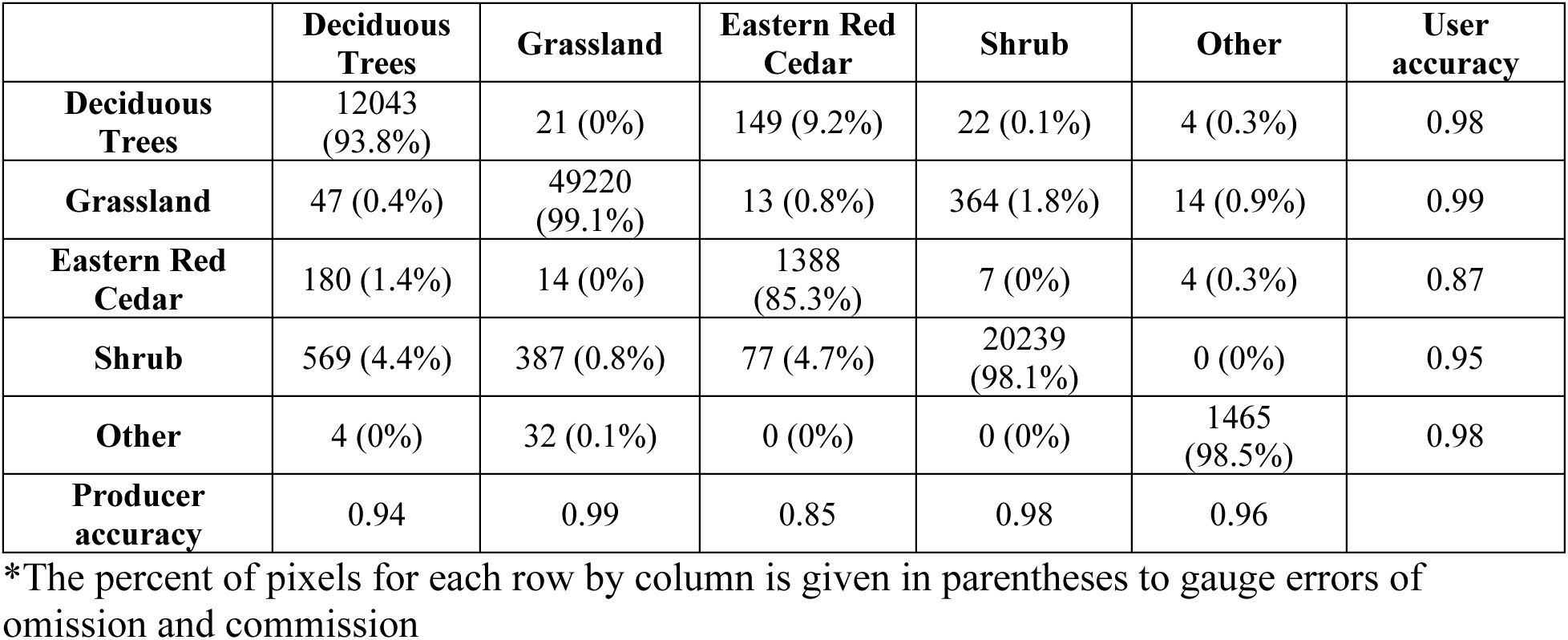
Confusion matrix for RF NAIP+NEON; columns represent class of evaluation pixels, and rows represent class of model classified pixels.

### Importance of different imagery data inputs

For NAIP, the most valuable input variable was red neighborhood (visualized in Fig. 3B), followed by blue neighborhood, red, and blue (Fig. 4a). The most valuable input variable for NEON was canopy height (visualized in Fig. 3C), followed far behind by NDVI and NDVI neighborhood (Fig. 4b). NAIP+NEON also largely relies on LiDAR, followed by five inputs from NAIP: red neighborhood, blue neighborhood, red localized, blue localized, and green neighborhood. In the combined model, NEON derived products (other than canopy height) provided very minimal GINI decrease, indicating weak contributions to increasing model accuracy (Fig 4C).

**Figure 4:**
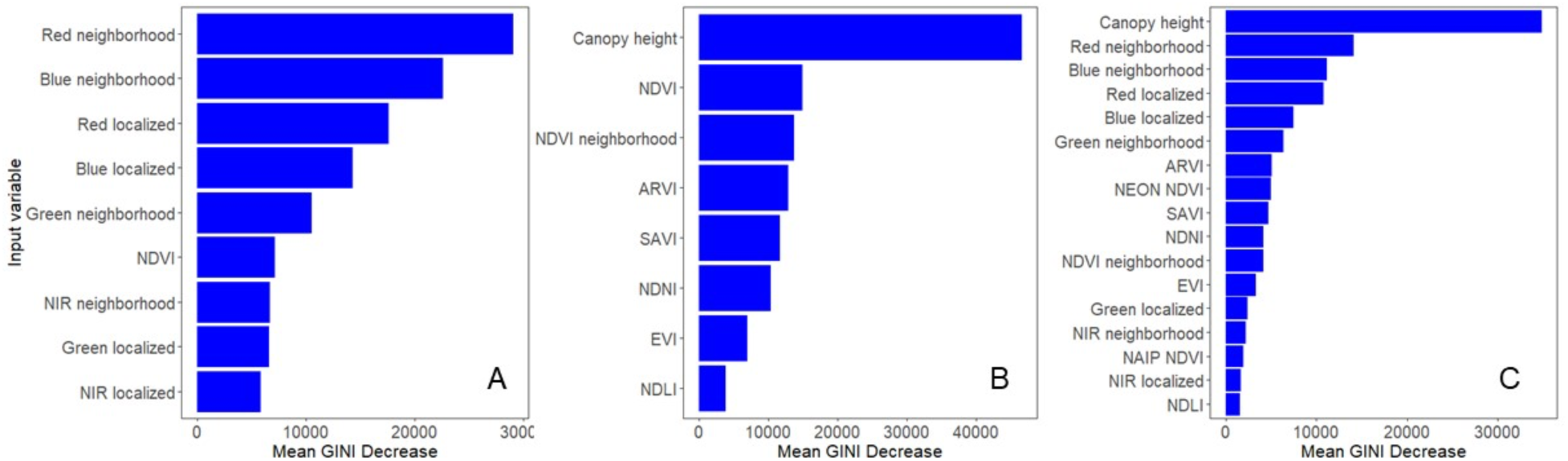
Average importance of each variable in Random Forest model building for: A) NAIP only; B) NEON only; and C) NAIP and NEON together. Mean GINI decrease essentially measures the amount of accuracy lost when that variable is removed. See Table 1 for input definitions.

### Model run time and other logistics

The amount of time it took to train each model on the training data varied greatly, with the shortest time at only 23 minutes for RF NEON, and the longest at 4 hours and 49 minutes for SVM NAIP (Table S2). RF and SVM models had substantial differences between training run times, with the slowest RF model taking 1 hour and 5 minutes, and the fastest SVM run time at 1 hour and 37 minutes (Table S2). The time it took the models to predict the entire study site (8,781,520 pixels) was drastically different between RFs and SVMs; RF classification took 7:20-10:43 minutes, whereas SVM classification took 2:05-6:15 hours (Table S3). For SVMs, model run time was negatively correlated with accuracy, with the NAIP+NEON model taking the least time to both train and classify, and being the most accurate (Tables 3, S2, & S3). However, for RF, the most accurate model (NAIP+NEON) took the longest time to train, but the shortest time to classify (Tables 3, S2, & S3).

## Discussion

Until recently, NAIP was the only widely available high-resolution open source of aerial remote sensing in the U.S.A. (Maxwell et al. 2017). We found that with a few manipulations (use of neighborhoods) and readily available machine learning methods, NAIP imagery succeeds at identifying grasslands and, to some extent, shrubs and deciduous trees. However, additional data, such as NEON, substantially increased our ability to correctly identify all forms of woody vegetation, especially ERC. The addition of NEON also made classification less subject to choices of machine learning methods. Therefore, in the locations where NEON is available (81 total sites vs entire continental U.S.A. for NAIP coverage), the two data sources could be used synergistically—NEON for its addition of LiDAR and NAIP for its undistorted red, green, blue, and NIR.

ERC is a native evergreen encroaching rapidly in tallgrass prairies, negatively impacting species diversity and ecosystem services (Briggs et al. 2002, Limb et al. 2010, Van Auken 2009, Zou et al. 2018). Detecting ERC is critical for monitoring and managing the impacts of woody encroachment (Meneguzzo and Liknes 2015). However, ERC had the lowest accuracy in all models, with PA lower than UA (Table 3), indicating that ERC is was undercounted, in some cases by almost half, because many ERC pixels are being classified as deciduous trees. This problem was acute when we only used NAIP imagery (Tables 4-5). We hypothesized that ERC would have much higher accuracy, since it has a much different leaf structure and water content than other woody plants in the area—characteristics that should be measured by NEON-derived products (e.g., NDVI, NDLI, NDNI). However, our hypothesis was incorrect, which has implications for using these models to make predictions of the rate and volume of current and future woody encroachment. Other studies also had difficulty detecting ERC in aerial imagery, specifically when ERC was at low densities (Kaskie et al. 2019, Kaskie et al. 2022). At densities below 30%, Kaskie et al. (2019) found <50% accuracy in ERC classification using aerial imagery alone. However, using other predictor variables such as slope, aspect, and Euclidean distance to the nearest ERC pixel, Kaskie et al. (2022) were able to increase the accuracy of ERC at low densities (<15%) to 84.7%. ERC density is low across our site, but accuracy was high (78-94%; Table 3) in models including NEON data. Therefore, NEON appears to overcome some challenges of low-density ERC detection and, presumably, other evergreen tree species. However, more work is needed to determine better and more efficient methods to accurately classify ERC at low densities.

We found that NEON imagery adds substantial accuracy when using machine learning methods to classify the remaining woody vegetation types (shrub and deciduous trees). Comparing the two single-source models (NEON vs NAIP), NEON increased accuracy by 8-32% for ERC, 6-24% for deciduous trees, and 5-13% for shrub cover (measured as percent increases going from NEON to NAIP, Table 3). In both models that included NEON data, canopy height data (based on NEON LiDAR), was by far the most influential input for accurately classifying our vegetation classes when using RF models (Fig. 4), indicating that LiDAR alone seems to be responsible for the increase in accuracy for woody plants. The importance of LiDAR was somewhat surprising given that all estimated values of canopy height <2 m are truncated to a value of zero as part of NEON’s data cleaning, and some areas dominated by woody vegetation on site are in the range of 1.5 to 2 m height. However, this is consistent with many other studies which also found that NEON LiDAR-based canopy height was among or the most important input for accurately classifying vegetation types, including separating woody plants from herbaceous plants and differentiating between species of tree species (Scholl et al. 2020, Pervin et al. 2022) and that LiDAR increases classification accuracy (Bork and Su 2007, Jin and Mountrakis 2022). Jin and Mountrakis (2022) analyzed 37 studies which compared accuracy using multispectral imagery alone versus classification with LiDAR (non-NEON sources) added and found increases in model accuracy for almost all studies, with accuracy increasing as much as 68%. This again points towards value added from NEON, but specifically from the addition of LiDAR, as NEON vegetation indices other than canopy height played a minor role in increasing model accuracy in the NAIP+NEON model (Fig. 4). NEON also provides a full suite of hyperspectral data, which were not tested here, but could have yet more value added on top of NEON’s LiDAR product. The hyperspectral data, however, is several hundred gigabytes large, and using it would not be consistent with our goal to create a method that is accessible to people of all skill levels and average computing resources.

Shrubs and small trees have been difficult to classify in the past because they are often smaller than the resolution of remotely sensed aerial imagery (> 9 m^2^; Whiteman and Brown 1998), leading to undercounting of shrubs and small trees (Brandt et al. 2020). Thus, we had hypothesized that shrubs would be among our lowest accuracy by class. However, we found high accuracy for shrubs in all models, but especially for NEON and NAIP+NEON models (94-98% vs. 82-93% for NAIP only; Table 3). Our other results show increased accuracy linked to the addition of LiDAR-based canopy height. However, the LiDAR data truncates all values below 2 m to 0 yet some shrubs are below 2 m tall. The accurate classification for shrubs indicate that another NEON remote sensing input is responsible for the increase in shrub accuracy. Some of this success may be because one of the encroaching species—*C. drummondii*—can produce high leaf area index values, with taller shrubs reaching values typical of dense deciduous or tropical forests (Tooley et al. 2024). Therefore, it remains unclear if the data sources and methods we used would perform as well in grasslands with WPE by shorter less dense species.

Finally, we found that two commonly used machine learning techniques performed similarly. RF models added a small amount of accuracy (< 2.6%), but the bigger difference was that RFs ran substantially faster than SVM models in program R (Table S2 & S3); prediction times for RF took minutes while SVM took hours. While the differences in run time meant that most models could train a machine learning model and project it to our study site in one day, run time would become an issue at larger spatial extents and over multiple time periods. RF was also slightly less sensitive to the data source, whereas SVMs performed poorly using only NAIP imagery. Finally, RFs also have the advantage of easily parsing the importance of different input variables, which could again, be valuable for using multiple remote sensing platforms synergistically.

### Remote sensing platforms and democratizing remote sensing

Remote sensing now has many options, such as UAVs, planes, and a growing suite of satellite products. Others have already reviewed the relative advantages of each approach (e.g., Matese et al. 2015, Tang et al. 2015, Toth et al. 2016, Bansod et al. 2017, Fassnacht et al. 2024). Generally, the advantages of low-flying planes fall between these UAVs and satellites, with a greater spatial extent than most UAVs and finer spatial resolution (grain) than most satellites. Low-flying planes can also carry heavy payloads, such as hyperspectral cameras and high-return high-resolution LiDAR (e.g., Asner et al. 2012, Baldeck et al. 2015). The typical disadvantage of remote sensing via planes is the initial learning curve and expense of piloting planes, maintaining cameras and LiDAR under flight conditions, and the data cleaning/carpentry of the raw data. These limitations also mean remote sensing from low-flying planes often has less temporal resolution. For instance, NAIP images are typically available every two years (Maxwell et al. 2017), whereas user-based drones and most satellites provide multiple measurements per growing season (Matese et al. 2015, Toth 2016).

Publicly available platforms have negated some disadvantages of low-flying planes by concentrating expertise. Experienced pilots and protogrammars capture the data. Experienced geographers then perform the initial data cleaning and formatting. The result is data in a format familiar to most end-users (e.g., a raster in ‘.tif’ format) and free of distortions such as cloud cover, significant spectral changes due to weather, and seamlines. Specialization reduces many of the logistical and financial barriers to using high-resolution data for image classification—a key part of the stated goals of state and federal programs to democratize scientific pursuit (Maxwell et al. 2017, Nagy et al. 2021). At a minimum, our results suggest that publicly available imagery can provide proof of concept to motivate more involved approaches such as UAVs or proprietary satellites. And in some ecosystems, such as Flint Hills Tallgrass prairie, our results suggest that these publicly available imagery can be used to track vegetation transitions, such as WPE. Similarly, contemporary and historical aerial imagery from planes have been used to remote sense woody vegetation (Baldeck et al. 2015, Faasnacht et al. 2RS024, Rosen et al. 2024, Weinstein et al. 2024) and WPE before (e.g. Laliberte et al. 2004, Strand et al. 2007, Weisburg et al. 2007, Smith et al. 2008, Miller et al. 2017, Keen et al. 2022, Soubry and Guo 2022). The difference is that most of these studies using plane-based data were from ad-hoc imagery, whereas one of the sources we use here is available nationwide in the U.S. (USDA NAIP, Maxwell et al. 2017) and the other is available for at least one site for each of the most widespread terrestrial ecosystem types in North America (NEON, Nagy et al. 2021). Our study is also in the relative minority of attempting and succeeding at high-resolution remote sensing shrubs in high productivity ecosystems (but see LaLiberte et al. 2004, Strand et al. 2007, Soubry and Guo 2022).

## Conclusions

Management of woody plant encroachment, including clonal shrublands and ERC trees is costly in time, money, and effort (Bidwell et al. 2002, Morford et al. 2022). Timely detection of small individual shrubs and trees can allow managers to engage in preventative management while woody plants are still at low densities and less resistant to disturbances, such as fire. Tools like NEON’s LiDAR increases accuracy of ERC classification and accuracy of detecting other woody plants, which can help implement management interventions and identify areas of elevated wildfire risk. Furthermore, increase in all class accuracies when using NEON creates more accurate overall vegetation mosaics, allowing other users to use these classified maps for applications including hydrology (Keen et al. 2022) and habitat use (Silber et al. 2025).

