## Supplementary material for "Combining machine learning and publicly available aerial data (NAIP and NEON) to achieve high-resolution remote sensing of grass-shrub-tree mosaics in the Central Great Plains (U.S.A.)": Supplimentary materials

**Title:**

**Authors and their information:**

Brynn Noble[1] and Zak Ratajczak[1*]

[1]Division of Biology, Kansas State University, Manhattan, KS 66502

116 Ackert Hall
Manhattan KS, 66506, U.S.A.

**Open research statement:**

Data sets utilized for this research are as follows:

Noble, B. and Z. Ratajczak. 2022. WPE01 Assessing the value added of NEON for using machine learning to quantify vegetation mosaics and woody plant encroachment at Konza Prairie ver 1. Environmental Data Initiative. https://doi.org/10.6073/pasta/a7b40e41080460bb1123dcc7b6d4d942 (Accessed 2022-12-08). https://portal.edirepository.org/nis/mapbrowse?scope=knb-lter-knz&identifier=167&revision=1

Code is provided as private-for-peer review at Figshare via the following link: https://figshare.com/s/9d0f3ae0f13385109551 Upon acceptance, code will be provided via Figshare.

Table S1: Best SVM model inputs calculated in R. Kernel refers to the method of data transformation. Degree is the degree of the polynomial kernel function. Gamma is the kernel coefficient which determines how much curvature will be in the decision boundary. Cost helps control error, a lower cost will accept a lower number of misclassified pixels.

| Image | Kernel | Degree | Gamma | Cost |
|---|---|---|---|---|
| NAIP | Radial | 3 | 0.1111 | 1 |
| NEON | Radial | 3 | 0.125 | 1 |
| NAIP+NEON | Radial | 3 | 0.0588 | 1 |

Table S2: Time to train models (300,328 pixels)

| **Model Run Time** | NAIP | NEON | NAIP+NEON |
|---|---|---|---|
| SVM | 4:49:00 | 1:43:00 | 1:37:00 |
| RF | 0:30:00 | 0:23:00 | 1:05:00 |

Table S3: Time to use models to predict (classify) entire study site (8,781,520 pixels).

| **Model Run Time** | NAIP | NEON | NAIP+NEON |
|---|---|---|---|
| SVM | 6:15:25 | 3:05:08 | 2:04:59 |
| RF | 0:09:59 | 0:10:43 | 0:07:20 |
